# Atomic context-conditioned protein sequence design using LigandMPNN

**DOI:** 10.1101/2023.12.22.573103

**Authors:** Justas Dauparas, Gyu Rie Lee, Robert Pecoraro, Linna An, Ivan Anishchenko, Cameron Glasscock, D. Baker

## Abstract

Protein sequence design in the context of small molecules, nucleotides, and metals is critical to enzyme and small molecule binder and sensor design, but current state-of-the-art deep learning-based sequence design methods are unable to model non-protein atoms and molecules. Here, we describe a deep learning-based protein sequence design method called LigandMPNN that explicitly models all non-protein components of biomolecular systems. LigandMPNN significantly outperforms Rosetta and ProteinMPNN on native backbone sequence recovery for residues interacting with small molecules (63.3% vs. 50.4% & 50.5%), nucleotides (50.5% vs. 35.2% & 34.0%), and metals (77.5% vs. 36.0% & 40.6%). LigandMPNN generates not only sequences but also sidechain conformations to allow detailed evaluation of binding interactions. Experimental characterization demonstrates that LigandMPNN can generate small molecule and DNA-binding proteins with high affinity and specificity.

**One-sentence summary:** We present a deep learning-based protein sequence design method that allows explicit modeling of small molecule, nucleotide, metal, and other atomic contexts.

## Main text

Protein sequence design methods seek to design, given a protein backbone structure, a sequence of amino acids that will fold to this backbone. Both physically based methods such as Rosetta (*1*) and deep-learning-based models such as ProteinMPNN (*2*), IF-ESM (*3*), and others (*4–9*) have been developed to solve this problem. The deep learning-based methods outperform physically based methods in designing sequences for protein backbones but currently available models can not incorporate non-protein atoms and molecules. For example, ProteinMPNN only explicitly considers protein backbone coordinates while ignoring any other atomic context. This context is critical for designing enzymes (*10*), protein-DNA/RNA (*11*), protein-small molecule (*12–14*), and protein-metal binders (*15*).

To enable the design of this wide range of protein functions, we set out to develop a deep learning method for protein sequence design that explicitly models the full non-protein atomic context. We sought to do this by generalizing the ProteinMPNN architecture to incorporate non-protein atoms. As with ProteinMPNN, we treat protein residues as nodes and introduce nearest neighbor edges based on Cα-Cα distances to define a sparse protein graph (Figure 1); protein backbone geometry is encoded into graph edges through pairwise distances between N, Cα, C, O, and virtual Cβ atoms. These input features are then processed using three encoder layers with 128 hidden dimensions to obtain intermediate node/edge representations. We experimented with introducing two additional protein-ligand encoder layers to encode protein-ligand interactions. We reasoned that only the nearest ligand atoms in 3D space would affect protein residue identities and side-chain conformations when protein backbone and ligand atoms were fixed.

**Fig. 1.**
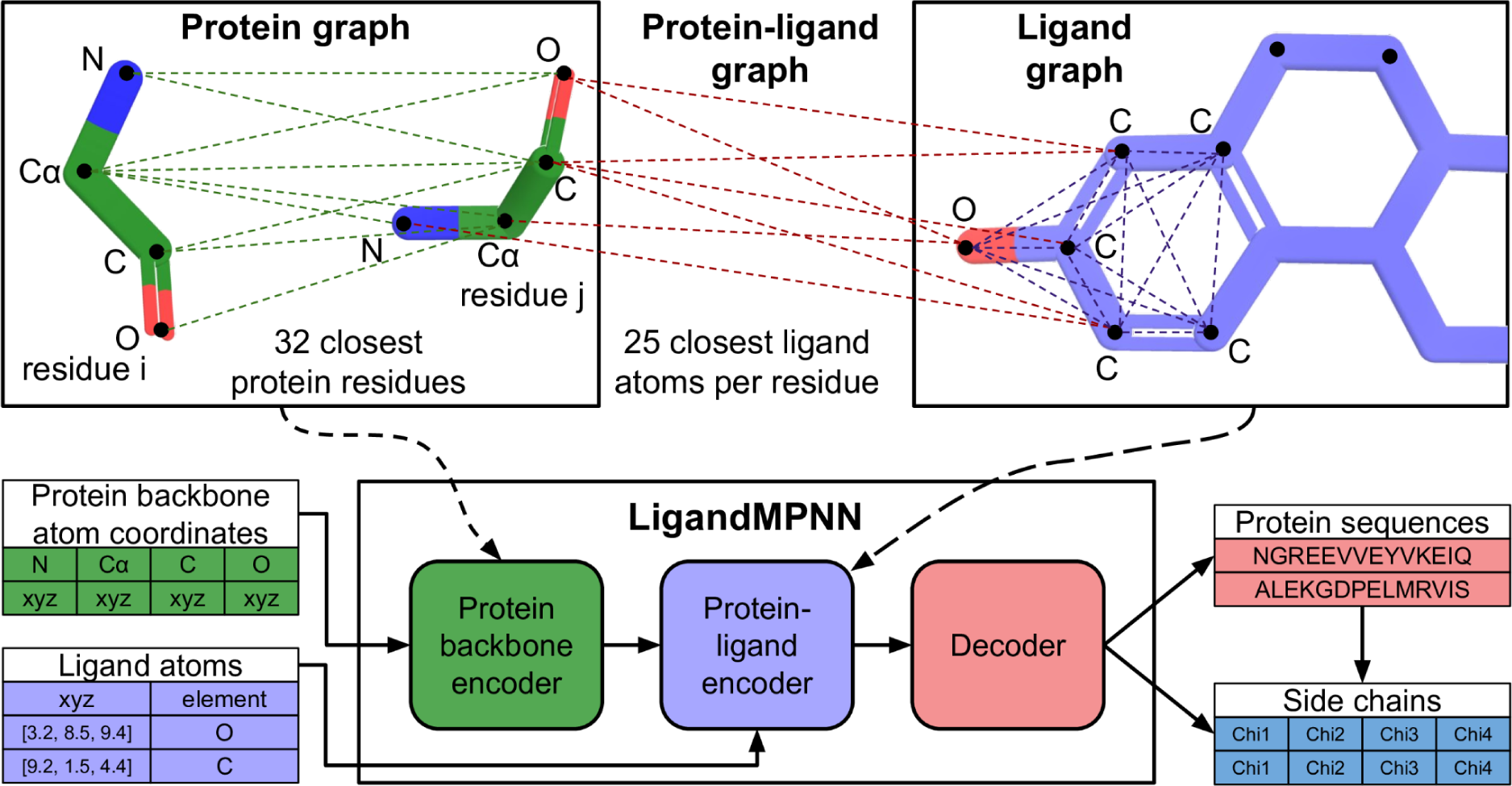
The LigandMPNN model. LigandMPNN operates on three different graphs. First, a protein-only graph with residues as nodes and 25 distances between N, Cα, C, O, and virtual Cβ atoms for residues i and j. Second, an intra-ligand graph with atoms as nodes that encodes chemical element types and distances between atoms as edges. Third, a protein-ligand graph with residues and ligand atoms as nodes and edges encoding residue j and ligand atom geometry. The LigandMPNN model has three neural network blocks: a protein backbone encoder, a protein-ligand encoder, and a decoder. Protein sequences and side-chain torsion angles are autoregressively decoded to obtain sequence and full protein structure samples. Metaparameter variation and ablation experiments are described in Supp. Fig. S1A-E.

To transfer information from ligand atoms to protein residues, we construct a protein-ligand graph with protein residues and ligand atoms as nodes and edges between each protein residue and the closest ligand atoms. We also build a fully connected ligand graph for each protein residue with its nearest neighbor ligand atoms as nodes; message passing between ligand atoms increases the richness of the information transferred to the protein through the ligand-protein edges. We obtained the best performance by selecting for the protein-ligand and individual residue intra-ligand graphs the 25 closest ligand atoms based on protein virtual Cβ and ligand atom distances (Figure S1A). The ligand graph nodes are initialized to one-hot encoded chemical element types, and the ligand graph edges to the distances between the atoms (Figure 1). The protein-ligand graph edges encode distances between N, Cα, C, O, and virtual Cβ atoms and ligand atoms (Figure 1). The protein-ligand encoder consisted of two message-passing blocks that updated the ligand graph representations and then updated the protein-ligand graph representations. The output of the protein-ligand encoder is combined with the protein encoder node representations and passed into the decoder layers. We call this combined protein-ligand sequence design model LigandMPNN.

To facilitate the design of symmetric (*15–16*) and multi-state proteins (*17*), we use a random autoregressive decoding scheme to decode the amino acid sequence as in the case of ProteinMPNN. With the addition of the ligand atom geometry encoding and the extra two protein-ligand encoder layers, the LigandMPNN neural network has 2.62 million parameters compared with 1.66 million ProteinMPNN parameters. Both networks are high-speed and lightweight (ProteinMPNN - 0.6 seconds and LigandMPNN - 0.9 seconds on a single CPU for 100 residues), scaling linearly with respect to the protein length. We augmented the training dataset by randomly selecting a small fraction of protein residues (2-4%) and using their side-chain atoms as context ligand atoms in addition to any small molecule, nucleotide, and metal context. While this augmentation did not significantly increase sequence recoveries (Figure S1B), training in this way also enables the direct input of side-chain atom coordinates to LigandMPNN to stabilize functional sites of interest.

We trained a sidechain packing neural network using the basic LigandMPNN architecture to predict the four side-chain torsion angles for each residue given a protein backbone and amino acid sequence inputs. Instead of predicting categorical distribution for amino acids, we predict a mixture (three components) of circular normal distributions for torsion angles chi1, chi2, chi3, and chi4: for each residue, we predict three mixing coefficients, three means, and three variances per chi angle. We autoregressively decompose the joint chi angle distribution by decoding all chi1 angles first, then all chi2 angles, chi3 angles, and finally all chi4 angles (After the model decodes one of the chi angles, its angular value, and the associated 3D atom coordinates are used for further decoding).

LigandMPNN was trained on protein assemblies in the PDB (as of Dec 16, 2022) determined by X-ray crystallography or cryoEM to better than 3.5 Å resolution and with a total length of less than 6,000 residues. The train/test split was based on protein sequences clustered at a 30% sequence identity cutoff. We evaluated LigandMPNN sequence design performance on a test set of 317 protein structures containing a small molecule, 74 with nucleic acids, and 83 with a transition metal (Figure 2A). For comparison, we retrained ProteinMPNN on the same training dataset of PDB biounits as LigandMPNN, except none of the context atoms were provided during training. Protein and context atoms were noised by adding 0.1 Å standard deviation Gaussian noise to avoid protein backbone memorization (*2*). We report native sequence recovery for residues that are close to the ligand, having side-chain atoms within 5.0 Å of any non-protein atoms. The median sequence recoveries (ten designed sequences per protein) near small molecules were 50.4% for Rosetta using the genpot energy function (*18*), 50.4% for ProteinMPNN, and 63.3% for LigandMPNN. For residues near nucleotides, median sequence recoveries were 35.2% for Rosetta (*11*) (using Rosetta energy optimized for protein-DNA interfaces), 34.0% for ProteinMPNN, and 50.5% for LigandMPNN, and for residues near metals, 36.0% for Rosetta (*18*), 40.6% for ProteinMPNN, and 77.5% for LigandMPNN (Figure 2A). Sequence recoveries were consistently higher for LigandMPNN over most proteins in the validation data set (Figure 2B; performance was correlated likely reflecting variation in the crystal structure and the amino acid composition of the site). LigandMPNN predicts amino acid probability distributions and uncertainties for each residue position: the expected confidence correlates with the actual sequence recovery accuracy (Figure 3C).

**Fig 2.**
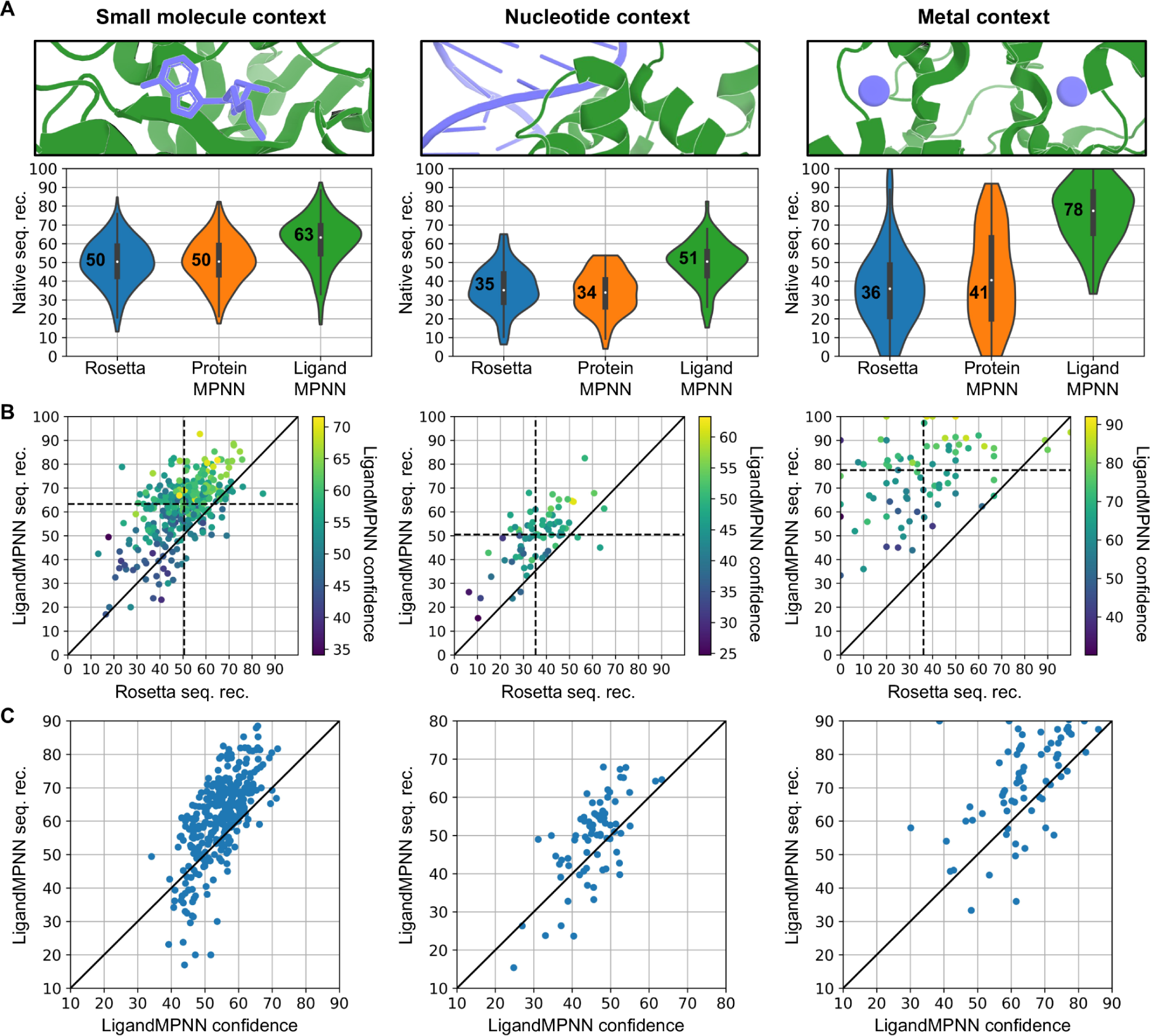
In silico evaluation of LigandMPNN sequence design. (**A**) LigandMPNN has a higher recovery of native protein sequences than Rosetta and ProteinMPNN around small molecules, nucleic acids, and metals. Sequence recoveries are averaged over the residues within 5.0 Å from the context atoms. (**B**) LigandMPNN has higher sequence recovery around non-protein molecules than Rosetta for most proteins. The color indicates LigandMPNN predicted confidence (between 0 and 100) about that protein. (**C**) Native sequence recovery correlates with LigandMPNN predicted confidence for designed sequences. One dot represents an average sequence recovery over ten sequences for one protein.

**Fig 3.**
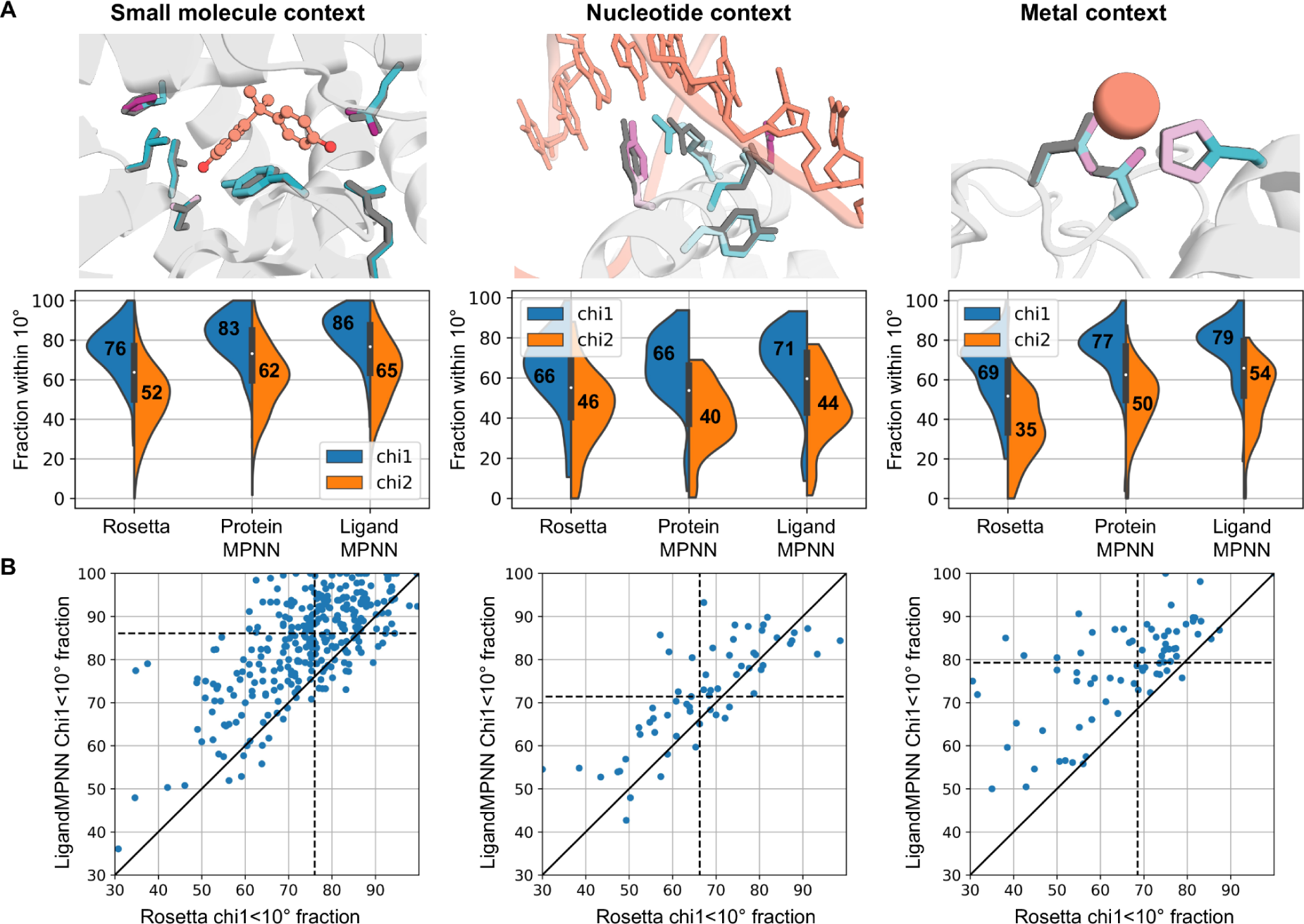
Evaluation of LigandMPNN side-chain packing accuracy. (**A**) Comparison of crystal side-chain packing (gray) with LigandMPNN side-chain packing (colored side chains by model confidence, teal is high, purple is low confidence per chi angle) for 2P7G, 1BC8, 1E4M proteins. The context atoms are shown in orange (small molecule, DNA, zinc). LigandMPNN has higher chi1 & chi2 torsion angle recovery (fraction of residues within 10° from native) than Rosetta and ProteinMPNN. (**B**) Per protein comparison of chi1 fraction recovery for LigandMPNN vs. Rosetta. One dot represents an average chi1 recovery over ten side-chain packing samples for one protein.

To assess the contributions to this highly sustained performance, we evaluated versions in which meta parameters and features were varied or ablated (Fig. S1A-E.) Decreasing the number of context atoms per residue primarily diminished sequence recovery around nucleic acids, likely because these are larger and contain more atoms than small molecules and metals (Fig. S1A). Providing side-chain atoms as additional context does not significantly affect LigandMPNN performance (Fig. S1B). As observed for ProteinMPNN, sequence recovery is inversely proportional to the amount of Gaussian noise added to input coordinates. The baseline model was trained with 0.1 Å standard deviation noise to reduce the extent to which the native amino acid can be read out simply based on the local geometry of the residue – crystal structure refinement programs introduce some memory of the native sequence into the local backbone. Training with 0.05 Å and 0.2 Å noise instead increased and decreased sequence recovery by about 2%, respectively (Fig. S1C; when comparing performance across methods, similar levels of noising must be used). Ablating the Protein-Ligand and Ligand graphs led to a 3% decrease in sequence recovery (Fig. S1D). Training on side-chain context atoms only (no small molecules, nucleotides, or metals) reduced sequence recovery around small molecules by 3.3% (Fig. S1E). Finally, a model trained without chemical element types as input features had much lower sequence recovery near metals (8% difference, Fig. S1D), but almost the same sequence recovery near small molecules and nucleic acids suggesting that the model can to some extent infer chemical element identity from bonded geometry.

We evaluated LigandMPNN side-chain packing performance on the same dataset for residues within 5.0 Å from the context atoms. We generated ten side-chain packing examples with the fixed backbone and fixed ligand context using Rosetta, LigandMPNN, and LigandMPNN without ligand context (called ProteinMPNN in Figure 3). The median chi1 fraction (within 10° from crystal packing) near small molecules was 76.0% for Rosetta, 83.3% for ProteinMPNN, and 86.1% for LigandMPNN, near nucleotides 66.2%, 65.6%, and 71.4%, and near metals 68.6%, 76.7%, and 79.3% for the three models (Figure 3A). LigandMPNN has a higher chi1 fraction recovery compared with Rosetta on most of the test proteins (Figure 3B), but only marginally better than ProteinMPNN (Figure S3C), suggesting that most of the information about side-chain packing is coming from the protein context rather than from the ligand context, consistent with binding site pre-organization. All the models struggle to predict chi3 and chi4 angles correctly. For LigandMPNN, weighted average fractions of correctly predicted chi1, chi2, chi3, and chi4 angles for the small molecule dataset were 84.0%, 64.0%, 28.3%, 18.7%, for Rosetta 74.5%, 50.5%, 24.1%, 8.1%, and for ProteinMPNN 81.6%, 60.4%, 26.7%, 17.4% (Figure S3B). The side-chain root-mean-square deviations (RMSD) are similar between the different methods as shown in Figures S4 and S5. Comparing ProteinMPNN vs LigandMPNN the biggest improvements in terms of RMSD are obtained for glutamine (Q) in the small molecule dataset, for arginine (R) in the nucleotide dataset, and for histidine (H) in the metal context dataset (Fig. S5) consistent with the important roles of interactions of these residues with the corresponding ligands.

### Experimental validation

We tested the capability of LigandMPNN to design binding sites for small molecules starting from previously characterized designs generated using Rosetta that either bound weakly or not at all to their intended targets: the muscle relaxant rocuronium, for which no binding was previously observed (Fig 4A) and the primary bile acid cholic acid (Fig 4B) for which binding was very weak (*13*,*14*). LigandMPNN was used to generate sequences around the ligands using the backbone and ligand coordinates as input; these retain and/or introduce new sidechain-ligand hydrogen bonding interactions. LigandMPNN redesigns either rescued binding (Fig. 4A) or improved the binding affinity (Fig. 4B). As described in (14), starting from the crystal structure of a previously designed cholic acid-protein complex, LigandMPNN increased cholic acid binding affinity 100-fold.

**Fig 4.**
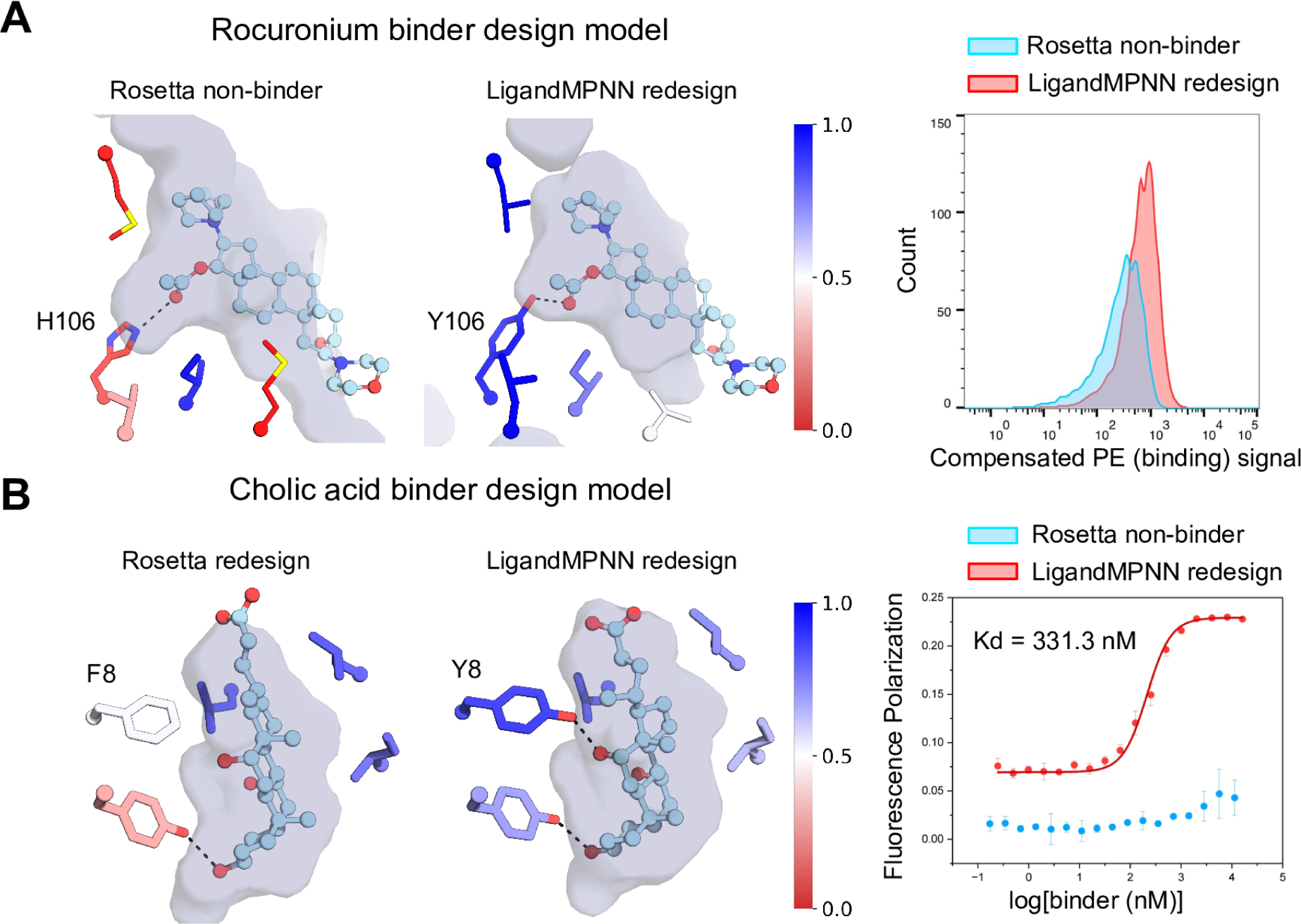
Rescue of Rosetta small molecule binder designs using LigandMPNN. (**A**) rocuronium, (**B**) cholic acid. Left panels: Interaction surface increases and the binding sites become more consistent with the target ligand (key binding residues are colored by LigandMPNN amino acid probabilities ranging from 0: red, 1: blue). Right panels, experimental data. (**A**) Flow cytometry of yeast displaying the designs following incubation with 1uM biotinylated rocuronium and SA-PE (**B**) Fluorescence polarization measurement of binding to cholic acid-FITC.

## Discussion

The deep learning-based LigandMPNN is superior to the physically based Rosetta for designing amino acids to interact with non-protein molecules. It is about 250 times faster (since the expensive Monte Carlo optimization over sidechain identities and compositions is completely bypassed), and the recoveries of native amino acid identities and conformations around ligands are consistently higher. The method is also easier to use since no expert customizations are required for new ligands (unlike Rosetta and other physically based methods which can require new energy function/force field parameters for new compounds). At the outset, we were unsure whether the accuracy of ProteinMPNN could extend to protein-ligand systems given the small amount of available training data, but our results suggest that for the vast majority of ligands, there is sufficient data. Nevertheless, we suggest some care in using LigandMPNN for designing binders to compounds containing elements occurring rarely or not at all in the PDB (in the latter case it is necessary to map to the most closely occurring element). Hybridization of the physically based and deep learning-based approaches may provide a better solution to the amino acid and sidechain optimization problems in the low data regime.

The accuracy of LigandMPNN has quickly made it the method of choice in our group for designing interactions of proteins with nucleic acids and small molecules, and these studies provide considerable additional experimental validation of the method. Glasscock et al. (*11*) developed a computational method for designing small sequence-specific DNA-binding proteins that recognize specific target sequences through interactions with bases in the major groove that uses LigandMPNN to design the protein-DNA interface. The crystal structure of a DNA binding protein designed with LigandMPNN recapitulated the design model closely (deposited to RCSB as PDB ID 8TAC). Lee et al. (*13*), An et al. (*14*), and Krishna et al. (12) used LigandMPNN to design small-molecule binding proteins with scaffolds generated by deep-learning and Rosetta-based methods. Iterative sequence design with LigandMPNN resulted in nanomolar to micromolar binders for the 17α-hydroxyprogesterone, apixaban, and SN-38 with NTF2-family scaffolds (13), nanomolar binders for cholic acid, methotrexate, and thyroxine (*14*) in pseudocyclic scaffolds, and binders for digoxigenin, heme, and bilin in RFdiffusion_allatom generated scaffolds (12). In total, more than 100 protein-DNA binding interfaces and protein-small molecule binding interfaces designed using LigandMPNN have been experimentally demonstrated to bind to their targets; this extensive experimental validation provides strong support for the power of the approach.

As with ProteinMPNN, we anticipate that LigandMPNN will be widely useful in protein design, enabling the creation of a new generation of small molecule binding proteins, sensors, and enzymes. To this end, we are making the code available at https://github.com/dauparas/LigandMPNN.

## Acknowledgments

We thank S. Pellock, Y. Kipnis, J. Wenckstern, A. Goncharenko, N. Hanikel, W. Ahern, P. Sturmfels, R. Krishna, D. Juergens, R. McHugh, P. Kim, and I. Kalvet for helpful discussions.

## Author contributions

Conceptualization: J.D., G.R.L, L.A., I.A.; Methodology: J.D., G.R.L, L.A., I.A, R.P., C.G..; Software: J.D., I.A..; Validation: G.R.L, L.A., R.P., C.G..; Formal analysis: J.D., G.R.L..; Resources: J.D., D.B.; Data curation: I.A., J.D., G.R.L, L.A., R.P.; Writing–original draft: J.D., D.B.; Writing–review and editing: J.D., D.B.; Visualization: J.D., G.R.L, L.A. Supervision: D.B..; Funding acquisition: J.D., D.B.

## Data and materials availability

All data are available in the main text or as supplementary materials. The LigandMPNN code is available at https://github.com/dauparas/LigandMPNN.

## Supplementary Materials for

### Materials and Methods

#### Methods for training LigandMPNN for sequence design

##### Training data

LigandMPNN was trained on a similar dataset to ProteinMPNN (*2*). We used protein assemblies in the PDB (as of Dec 16, 2022) determined by X-ray crystallography or cryoEM to better than 3.5 Å resolution and with less than 6,000 residues. We parsed all residues present in the PDBs except [“HOH”, “NA”, “CL”, “K”, “BR“]. Protein sequences were clustered at 30% sequence identity cutoff using mmseqs2 (*19*). We held out a non-overlapping subset of proteins that have small molecule contexts (a total of 317), nucleotide contexts (a total of 74), and metal contexts (a total of 83).

##### Optimizer and loss function

For optimization, we used Adam with beta1 = 0.9, beta2 = 0.98, and epsilon = 10−9 same as for ProteinMPNN. Models were trained with a batch size of 6k tokens, automatic mixed precision, and gradient checkpointing on a single NVIDIA A100 GPU for 300k optimizer steps. We used categorical cross entropy for the loss function following the ProteinMPNN paper (*2*).

##### Input featurization and model architecture

We used the same input features as in the ProteinMPNN paper for the protein part. For the atomic context input features, we used one-hot encoded chemical element types as node features for the ligand graph and the radial basis function encoded distances between the context atoms as edges for the ligand graph. To encode the interaction between protein-context atoms we used distances between N, Cα, C, O, and virtual Cβ atoms and context atoms. In addition, we added angle-based sin/cos features describing context atoms in the frame of N-Cα-C atoms.

We used the same MPNN architecture as used in ProteinMPNN for the encoder, decoder, and protein-ligand encoder blocks. Encoder and decoder blocks work on protein nodes and edges, i.e mapping vertices [N] and edges [N, K] to updated vertices [N] and edges [N, K] where N is the number of residues, and K is the number of direct neighbors per residue. We choose M context atoms per residue resulting in [N, M] protein-atom interactions. The ligand graph blocks map vertices of size [N, M] and edges of size [N, M, M] (fully connected context atoms) to updated vertices [N, M]. The updated [N, M] representation is used in the protein-ligand graph to map vertices [N] and edges [N, M] into updated vertices [N]. For more details refer to the LigandMPNN code.

##### Ablation studies

We trained several LigandMPNN variants to understand dependencies on hyperparameters (S1A-E). All sequence recoveries are reported with respect to the baseline model which was noised with 0.1 Å standard deviation, hidden dimension was 128, 3 encoder layers, 2 protein-context atom encoder layers, 2 context atom encoder layers, and 3 decoder layers, 25 context atoms per residue, 2% randomly sampled side-chains used as additional context atoms, all sequences were designed using the sampling temperature of 0.1. Firstly, we trained LigandMPNN with 2, 4, 8, 16, 25, 32 context atoms per residue (Fig S1A). We see that sequence recoveries increase with more context atoms given, but saturate at about 8-16 atoms. The most noticeable effect is on the nucleotide context residues. We also varied the fraction of side chains used as a context which were randomly chosen during training (Fig. S1B). There is a small increase in sequence recovery for nucleotide context, but more importantly, this allows the use of side-chain atoms during the inference for fixed activate site design examples. In practice, we do not know the exact backbone and context atom positions when designing sequences. Same as for ProteinMPNN we added Gaussian noise to all input coordinates to avoid backbone memorization. Sequence recovery goes down for the models trained with larger amounts of noise acting as regularization (Fig. S1C). For ablations (Fig. S1D), we trained models without Protein-Ligand and Ligand graphs by passing protein-context atom features flattened as vertex features into the encoder. We saw about a 3% reduction in sequence recovery. Removing just the Ligand graph and keeping the Protein-Ligand graph results in about a 1% decrease. Training without chemical element types has the most effect on the residues near metal ions. However, only a marginal difference for small molecule and nucleotide context suggests that the model can infer chemical elements from the geometry. Finally, we trained LigandMPNN only using protein side-chain atoms as context (Fig. S1E). Sequence recoveries near nucleotide and metal context went significantly down, however, we observed only about a 3-4% decrease in sequence recovery near small molecules, even though the model has never been trained on any small molecule data. This suggests that the model can generalize from side-chain to small-molecule contexts due to the similarity in carbon, oxygen, and nitrogen chemistry, and volume exclusion.

##### Sequence bias

For all three different datasets (small molecule, nucleotide, and metal) we plotted 1D (PSSM) and 2D (confusion matrix) sequence biases for Rosetta and LigandMPNN designs for the residues within 5.0 Å from the context atoms (Fig. S2A-C). We noticed that when LigandMPNN is uncertain about what amino acid to predict, it predicts the most likely amino acids in these contexts. For example, predicting a higher number of lysines near nucleotides and a slightly higher number of histidines near metals (Fig. S2B). We also noted that sequence recoveries are very much correlated between ProteinMPNN and LigandMPNN for small molecule and nucleotide contexts suggesting limited information coming from the backbone. There is almost no correlation near metal context suggesting that knowing the placement of metal ions provides very useful information to locally determine protein sequence (Fig. S2C).

#### Methods for training LigandMPNN for side-chain prediction

##### Side-chain modeling

We used the same architecture as for the sequence design to predict side-chain conformations. In this case, the model predicts a mixture (three components) of circular normal distributions for torsion angles chi1, chi2, chi3, and chi4. The objective was to maximize the log probability of training examples. We used the same training dataset as for sequence design with a 3.5 Å resolution cutoff. Training with higher-resolution cutoff (e.g. 2-2.5 Å) structures might improve high-accuracy side-chain modeling, but it does result in a much smaller number of nucleotide-protein structures available for training leading to worse performance for residue near different contexts. We trained a fully autoregressive model that factorizes a joint residue-chi angle distribution into conditional distributions. Since the chi1 angles have the lowest variance we autoregressively predict all chi1 angles first, and then autoregressively all chi2, chi3, and chi4 angles. The model predicts a chi angle distribution from which we sample a particular chi angle and pass its sin/cos values as well as 3D coordinates of corresponding atoms as an input for the next prediction. Figure S3 compares ProteinMPNN and Rosetta versus LigandMPNN using a 10° chi angle cutoff for residues within 5.0 Å from the context atoms. If chi1 is more than 10° away from the ground truth then chi2, chi3, and chi4 are considered to be incorrect. Figures S4 and S5 show RMSD deviations of side-chain atoms (all side-chain atoms excluding N, Cα, C, O, and Cβ).

**Fig S1.**
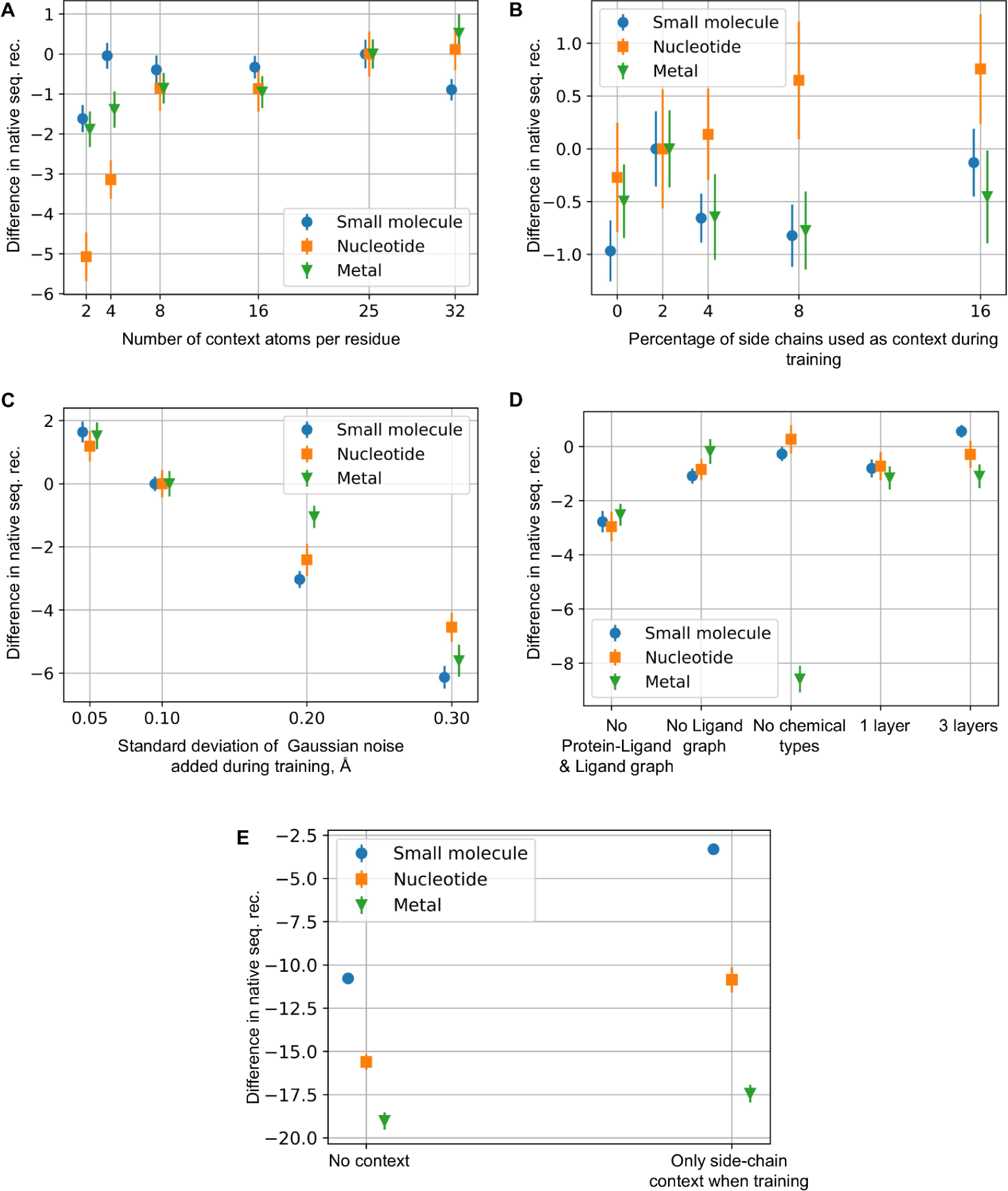
Different model variants. (**A**) The difference in native sequence recovery for LigandMPNN models trained with a different number of context atoms per residue. The baseline model was trained with 25 atoms per residue. (**B**) The difference in native sequence recovery as a function of the percentage of side chains used as context atoms during training. (**C**) Sensitivity of the sequence recovery to the magnitude of Gaussian noise added during training to protein & context atoms. (**D**) Sequence recovery change for different ablations. For the “No Protein-Ligand & Ligand graph,” chemical element types and geometry were used directly as node inputs for the protein graph. For the “No Ligand graph,” we kept 2 MPNN layers of Protein-Ligand only. For the “No chemical types,” we passed zeros as chemical element types when training and during the inference. For the “1/3 layers”, we used 1 or 3 instead of 2 baseline model Protein-Ligand & Ligand MPNN layers. (**E**) Comparing LigandMPNN with the model trained without any context (ProteinMPNN) and with the LigandMPNN model trained with only protein side-chain context.

**Fig S2.**
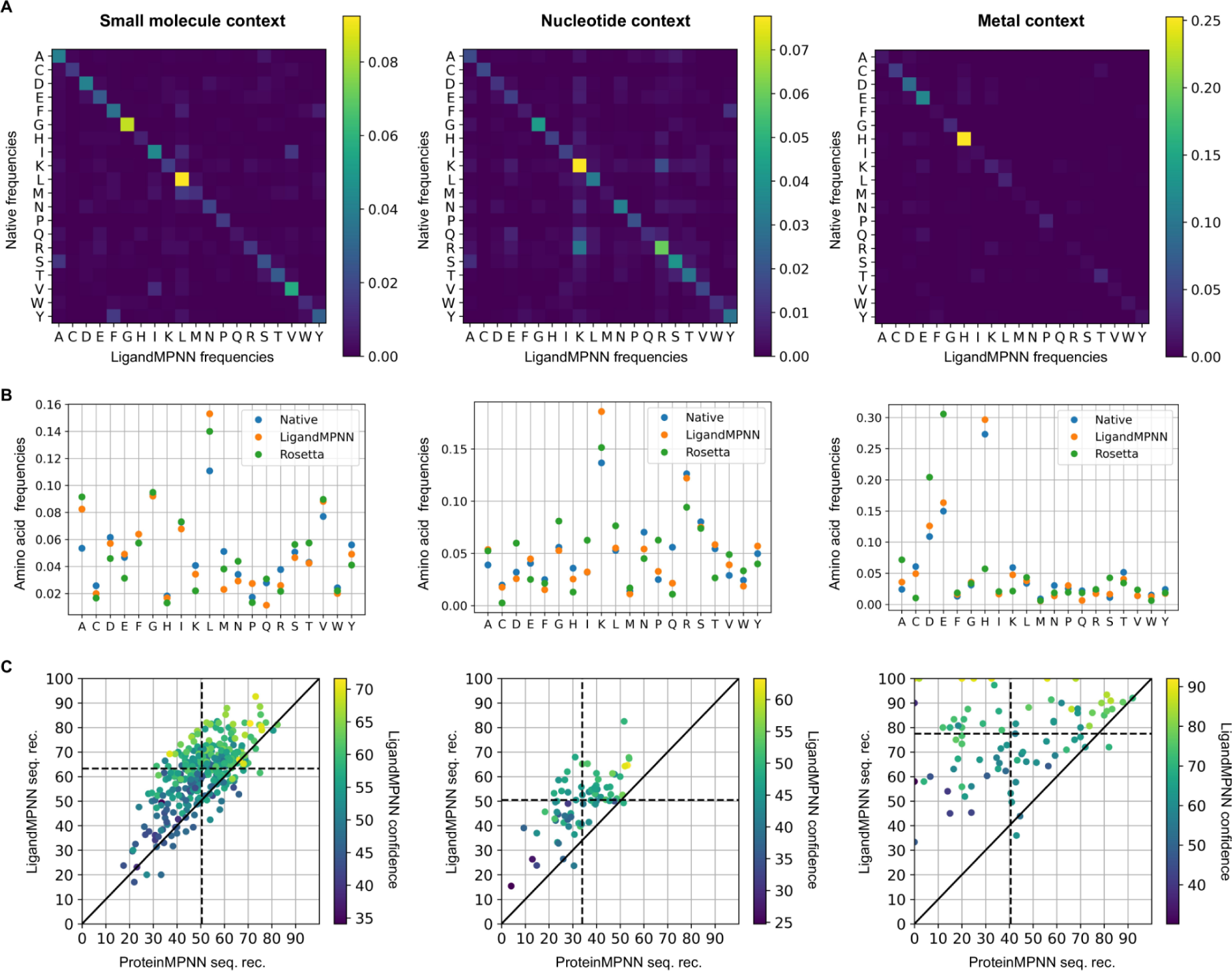
Sequence biases for residues within 5.0 Å from ligands. (**A**) Confusion matrices for LigandMPNN. (**B**) Amino acid biases comparing native vs LigandMPNN vs Rosetta sequences. (**C**) Comparing ProteinMPNN vs LigandMPNN per protein basis. Color represents LigandMPNN confidence.

**Fig S3.**
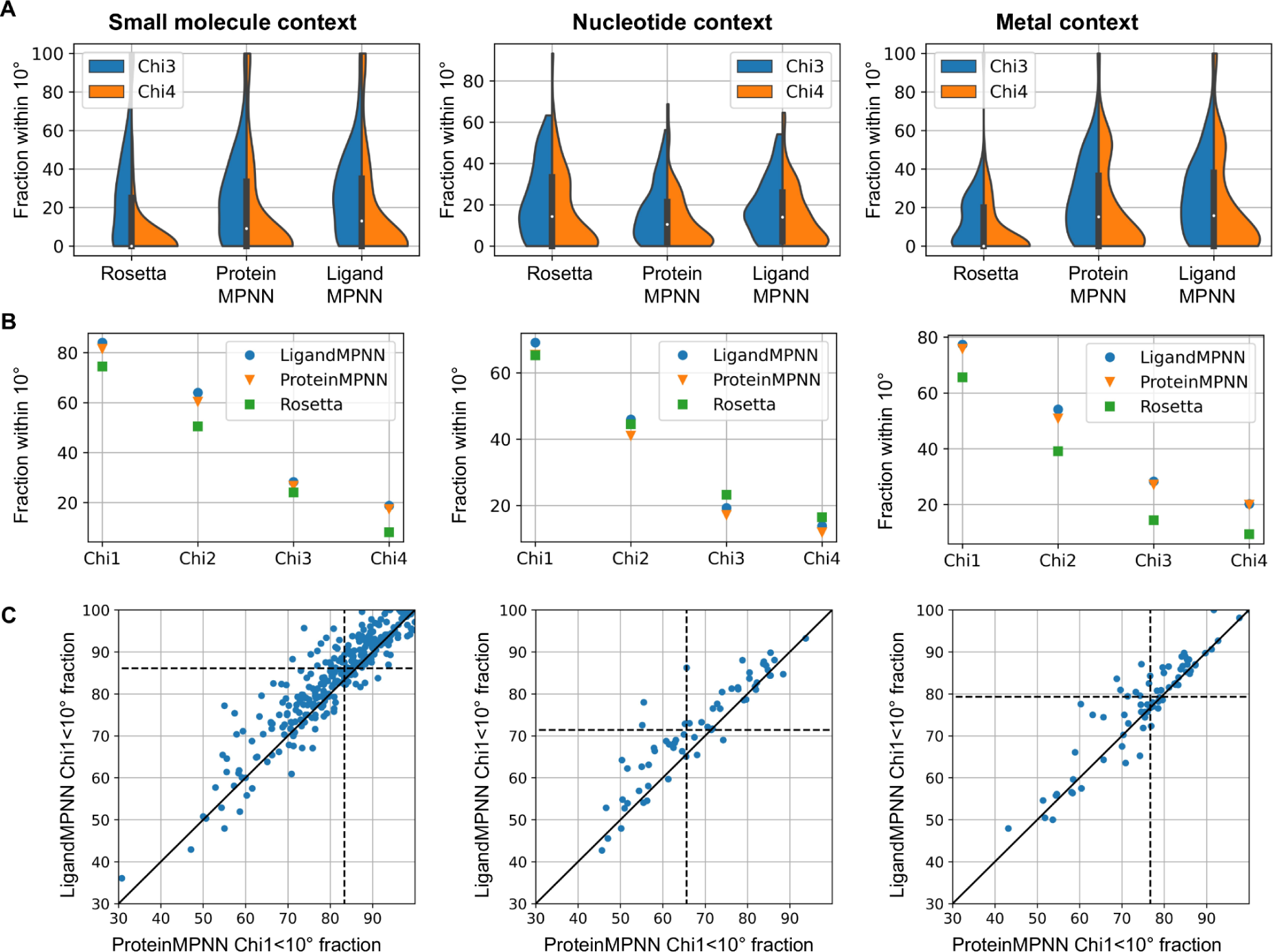
Side-chain chi angle recoveries for residues within 5.0 Å from ligands. (**A**) Shows distributions for fractions within 10 degrees from a native conformation of chi3 and chi4 for Rosetta, ProteinMPNN, and LigandMPNN. (**B**) Mean chi recoveries for Ligand mpn, ProteinMPNN, and Rosetta. (**C**) Per protein comparison of chi1 recovery of ProteinMPNN vs LigandMPNN.

**Fig S4.**
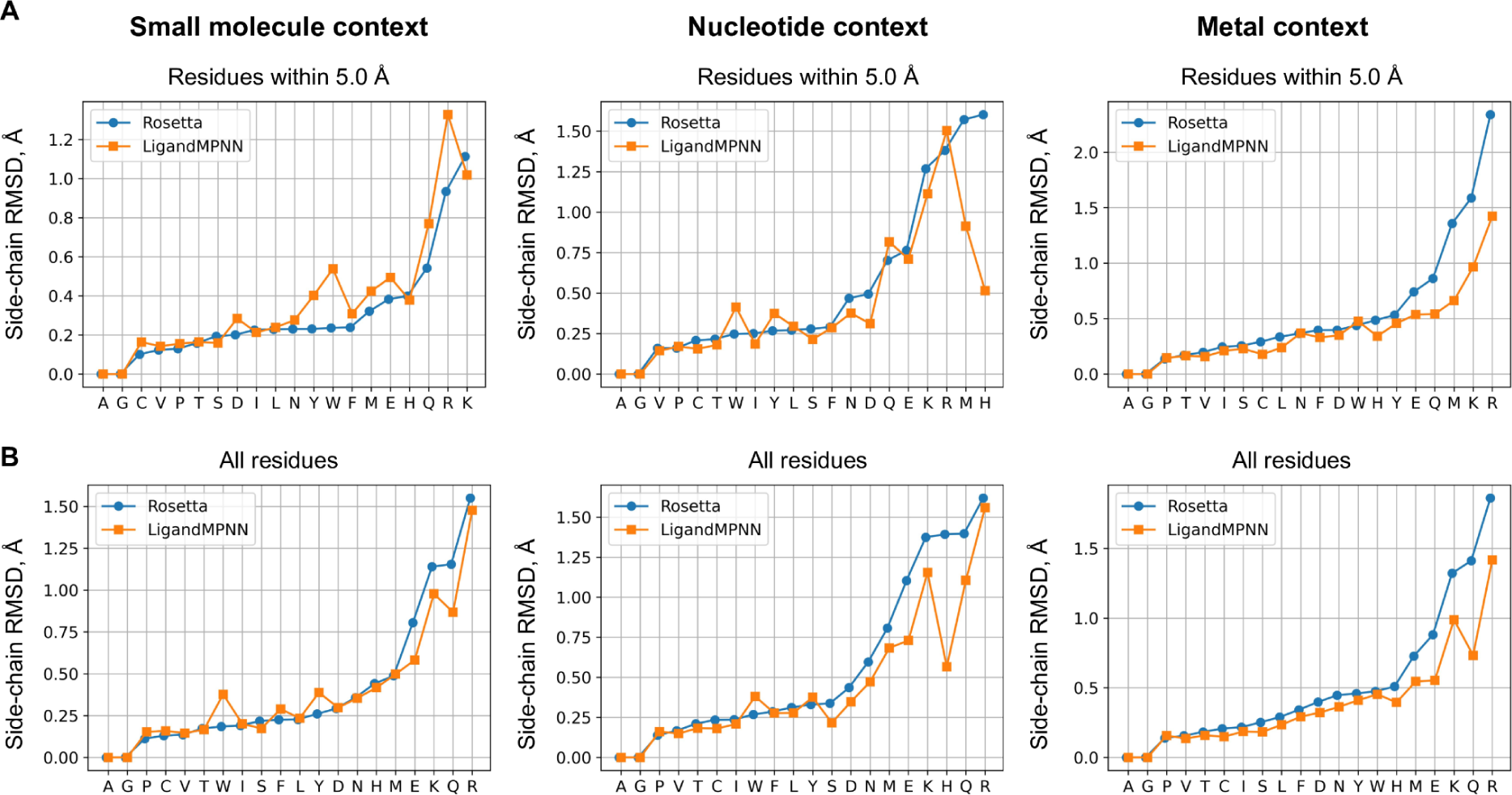
Median side-chain root-mean-square deviations (RMSD) per amino acid for Rosetta vs LigandMPNN. (**A**) RMSDs over protein residues within 5.0 Å from the context atoms, (**B**) over all residues in the protein chain.

**Fig S5.**
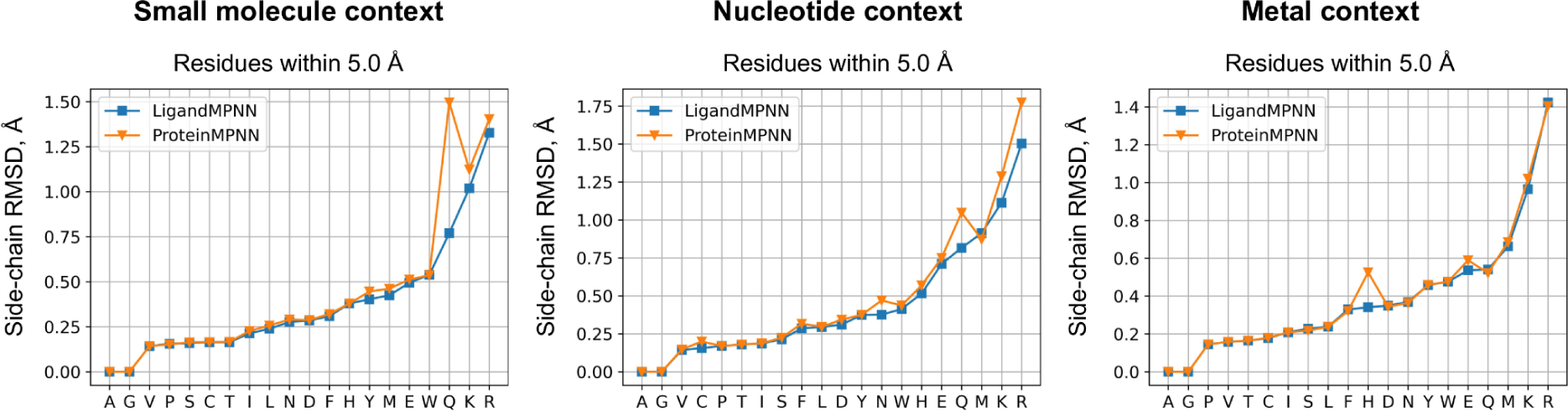
Comparing LigandMPNN vs ProteinMPNN for side-chain packing. The biggest improvements in terms of side-chain root-mean-square deviation (RMSD) are obtained for glutamine (Q) in the small molecule dataset, for arginine (R) in the nucleotide dataset, and for histidine (H) in the metal context dataset.

#### Small molecule context benchmark PDB IDs

*[’1a28’, ’1bzc’, ’1drv’, ’1e3g’, ’1elb’, ’1elc’, ’1epo’, ’1f0r’, ’1g7f’, ’1g7g’, ’1gvw’, ’1gx8’, ’1i37’, ’1kav’, ’1kdk’, ’1kv1’, ’1l8g’, ’1lhu’, ’1lpg’, ’1nc1’, ’1nfx’, ’1nhz’, ’1nl9’, ’1nny’, ’1nwl’, ’1ony’, ’1pyn’, ’1qb1’, ’1qkt’, ’1qxk’, ’1r0p’, ’1sj0’, ’1sqn’, ’1v2n’, ’1xjd’, ’1xws’, ’1yc1’, ’1yqj’, ’1z95’, ’1zp8’, ’2ayr’, ’2b07’, ’2b4l’, ’2baj’, ’2bak’, ’2bal’, ’2bsm’, ’2cet’, ’2e2r’, ’2f6t’, ’2fdp’, ’2g94’, ’2hah’, ’2ihq’, ’2iwx’, ’2j2u’, ’2j34’, ’2j4i’, ’2j94’, ’2j95’, ’2o0u’, ’2oax’, ’2ojg’, ’2ojj’, ’2p4j’, ’2p7g’, ’2p7z’, ’2pog’, ’2qbp’, ’2qbq’, ’2qbs’, ’2qe4’, ’2qmg’, ’2uwl’, ’2uwo’, ’2uwp’, ’2v7a’, ’2vh0’, ’2vh6’, ’2vkm’, ’2vrj’, ’2vw5’, ’2vwc’, ’2w8y’, ’2wc3’, ’2web’, ’2wec’, ’2weq’, ’2wgj’, ’2wuf’, ’2wyg’, ’2wyj’, ’2xab’, ’2xb8’, ’2xda’, ’2xht’, ’2xj1’, ’2xj2’, ’2xjg’, ’2xjx’, ’2y7x’, ’2y7z’, ’2y80’, ’2y81’, ’2y82’, ’2ydw’, ’2yek’, ’2yel’, ’2yfe’, ’2yfx’, ’2yge’, ’2ygf’, ’2yi0’, ’2yi7’, ’2yix’, ’2zmm’, ’3acw’, ’3acx’, ’3b5r’, ’3b65’, ’3bgq’, ’3bgz’, ’3ckp’, ’3cow’, ’3coy’, ’3coz’, ’3d7z’, ’3d83’, ’3eax’, ’3ekr’, ’3fv1’, ’3fv2’, ’3fvk’, ’3gba’, ’3gbb’, ’3gcs’, ’3gcu’, ’3gy3’, ’3hek’, ’3i25’, ’3ioc’, ’3iph’, ’3iw6’, ’3k97’, ’3lpi’, ’3lpk’, ’3lxk’, ’3m35’, ’3myg’, ’3n76’, ’3nq3’, ’3nyx’, ’3o5x’, ’3o8p’, ’3pww’, ’3roc’, ’3tfn’, ’3u81’, ’3ueu’, ’3uev’, ’3uew’, ’3uex’, ’3vha’, ’3vhc’, ’3vhd’, ’3vje’, ’3vvy’, ’3vw1’, ’3vw2’, ’3wha’, ’3wz6’, ’3wz8’, ’3zc5’, ’3zm9’, ’3zze’, ’4a4v’, ’4a4w’, ’4a7i’, ’4ag8’, ’4ap7’, ’4b6o’, ’4b9k’, ’4cd0’, ’4cga’, ’4cmo’, ’4da5’, ’4e5w’, ’4e6d’, ’4e9u’, ’4ea2’, ’4egk’, ’4er1’, ’4fcq’, ’4ffs’, ’4flp’, ’4g8n’, ’4gny’, ’4gu6’, ’4hge’, ’4igt’, ’4k0y’, ’4k9y’, ’4kao’, ’4kcx’, ’4lyw’, ’4m0r’, ’4m12’, ’4m13’, ’4muf’, ’4nh8’, ’4nwc’, ’4o04’, ’4o05’, ’4o07’, ’4o09’, ’4o0b’, ’4p5z’, ’4pmm’, ’4pop’, ’4qev’, ’4qew’, ’4qyy’, ’4rfm’, ’4rwj’, ’4twp’, ’4uyf’, ’4v01’, ’4w9f’, ’4w9l’, ’4wa9’, ’4wkn’, ’4×6p’, ’4xip’, ’4xir’, ’4y79’, ’4ybk’, ’4ymb’, ’4yml’, ’4ynb’, ’4yth’, ’4z0k’, ’4zae’, ’5aa9’, ’5acy’, ’5d26’, ’5d3h’, ’5d3j’, ’5d3l’, ’5d3t’, ’5dlx’, ’5dqc’, ’5dwr’, ’5e74’, ’5egm’, ’5eng’, ’5eqp’, ’5eqy’, ’5er1’, ’5exm’, ’5exn’, ’5f9b’, ’5fto’, ’5fut’, ’5hcv’, ’5i3v’, ’5i3y’, ’5i9x’, ’5i9z’, ’5ie1’, ’5ih9’, ’5jq5’, ’5kz0’, ’5l2s’, ’5lli’, ’5lny’, ’5lsg’, ’5neb’, ’5nw1’, ’5nyh’, ’5op5’, ’5oq8’, ’5qqp’, ’5t19’, ’5tpx’, ’5v82’, ’5yfs’, ’5yft’, ’6c2r’, ’6cjr’, ’6cpw’, ’6dgq’, ’6dgr’, ’6dyu’, ’6dyv’, ’6el5’, ’6elo’, ’6elp’, ’6ey9’, ’6eyb’, ’6f1n’, ’6ge7’, ’6gf9’, ’6gfs’, ’6ghh’, ’6i61’, ’6i64’, ’6i67’, ’6md0’, ’6mh1’, ’6mh7’, ’6n7a’, ’6n8x’, ’6no9’, ’6nv7’, ’6nv9’, ’6olx’, ’6qi7’]*

#### Nucleotide context benchmark PDB IDs

*[’1a0a’, ’1am9’, ’1an4’, ’1b01’, ’1bc7’, ’1bc8’, ’1di2’, ’1ec6’, ’1hlo’, ’1hlv’, ’1i3j’, ’1pvi’, ’1qum’, ’1sfu’, ’1u3e’, ’1xpx’, ’1yo5’, ’1zx4’, ’2c5r’, ’2c62’, ’2nq9’, ’2o4a’, ’2p5l’, ’2xdb’, ’2ypb’, ’2zhg’, ’2zio’, ’3adl’, ’3bsu’, ’3fc3’, ’3g73’, ’3gna’, ’3gx4’, ’3lsr’, ’3mj0’, ’3mva’, ’3n7q’, ’3olt’, ’3vok’, ’3vwb’, ’3zp5’, ’4ato’, ’4bhm’, ’4bqa’, ’4e0p’, ’4nid’, ’4wal’, ’5cm3’, ’5haw’, ’5mht’, ’5vc9’, ’5w9s’, ’5ybd’, ’6bjv’, ’6dnw’, ’6fqr’, ’6gdr’, ’6kbs’, ’6lff’, ’6lmj’, ’6od4’, ’6wdz’, ’6×70’, ’6y93’, ’7bca’, ’7c0g’, ’7el3’, ’7jsa’, ’7ju3’, ’7kii’, ’7kij’, ’7mtl’, ’7z0u’, ’8dwm’]*

#### Metal context benchmark PDB IDs

*[’1dwh’, ’1e4m’, ’1e6s’, ’1e72’, ’1f35’, ’1fee’, ’1job’, ’1lqk’, ’1m5e’, ’1m5f’, ’1moj’, ’1mxy’, ’1mxz’, ’1my1’, ’1nki’, ’1qum’, ’1sgf’, ’1t31’, ’1u3e’, ’2bdh’, ’2bx2’, ’2cfv’, ’2e6c’, ’2nq9’, ’2nqj’, ’2nz6’, ’2ou7’, ’2vxx’, ’2zwn’, ’3bvx’, ’3cv5’, ’3f4v’, ’3f5l’, ’3fgg’, ’3hg9’, ’3hkn’, ’3hkt’, ’3i9z’, ’3k7r’, ’3l24’, ’3l7t’, ’3m7p’, ’3mi9’, ’3o1u’, ’3u92’, ’3u93’, ’3u94’, ’3won’, ’4aoj’, ’4dy1’, ’4hzt’, ’4i0f’, ’4i0j’, ’4i0z’, ’4i11’, ’4i12’, ’4jd1’, ’4naz’, ’4wd8’, ’4×68’, ’5f55’, ’5f56’, ’5fgs’, ’5hez’, ’5i4j’, ’5l70’, ’5vde’, ’6a4x’, ’6buu’, ’6cyt’, ’6iv2’, ’6lkp’, ’6lrd’, ’6wdz’, ’6×75’, ’7dnr’, ’7e34’, ’7kii’, ’7n7g’, ’7s7l’, ’7s7m’, ’7w5e’, ’7wb2’]*

